# Real-time non-invasive intracranial state estimation using unscented Kalman filter

**DOI:** 10.1101/400945

**Authors:** Chanki Park, Seungjun Ryu, Bonghyun Jung, Sangpyong Lee, Changkie Hong, Yongbae Kim, Boreom Lee

## Abstract

Intracranial pressure (ICP) monitoring is desirable as a first-line measure to assist decision-making in cases of increased ICP. Clinically, non-invasive ICP monitoring is also required to avoid infection and hemorrhage in patients. The relationships among the arterial blood pressure (*P*_a_), ICP, cerebral blood flow, and its velocity (*Q*_CBFv_) measured by transcranial Doppler ultrasound measurement have been reported. However, real-time non-invasive ICP estimation using these modalities is less well documented. Here, we present a novel algorithm for real-time and non-invasive ICP monitoring with *Q*_CBFv_ and *P*_a_, called direct-current (DC)-ICP. This technique is compared with invasive ICP for 11 traumatic-brain-injury patients admitted to Cheju Halla Hospital and Gangnam Severance Hospital from July 2017 to June 2018. The inter-subject correlation coefficient between true and estimate was 0.70. The AUCs of the ROCs for prediction of increased ICP for the DC-ICP methods are 0.816. Thus, *Q*_CBFv_ monitoring can facilitate reliable real-time ICP tracking with our novel DC-ICP algorithm, which can provide valuable information under clinical conditions.

## Introduction

Invasive intracranial pressure (ICP) monitoring is desirable as a first-line measure to assist decision-making in cases of increased ICP, for those with a Glasgow coma scale (GCS) score < 8, and for those who need aggressive medical care^1^. Although the most accurate method is invasive monitoring, non-invasive ICP monitoring is also required to avoid infection and hemorrhaging under clinical conditions^2^.

Non-invasive ICP estimation would act as an aid for specific patients requiring invasive monitoring. For traumatic-brain-injury (TBI) patients, an ICP of less than 20−25 mmHg should be maintained^3^, with intensive care unit management for 7 days or more. However, if invasive ICP monitoring is prolonged for more than 5 days, infectious conditions develop in 85% of TBI patients^2^. Further, non-invasive monitoring may assist proper management of coagulopathy patients for whom invasive ICP measurement is not immediately available because the hemorrhagic risk is too high^4^.

Therefore, as an alternative to invasive ICP monitoring, non-invasive ICP monitoring devices that can accurately and continuously estimate ICP should be developed. Various methods of non-invasive ICP estimation have been studied to date, being based on measurement of related physiological variables such as the optic nerve sheath diameter^5^, phase-contrast magnetic resonance imaging of the blood and cerebrospinal fluid (CSF) flow^6^, electroenchephalogram signals of the visual evoked potentials^7^, and measurement of the tympanic membrane displacement^8^. However, these non-invasive methods require calibration for ICP estimation and cannot monitor ICP continuously.

For real-time ICP estimation, correlations between the ICP and *Q*_CBFv_ using transcranial Doppler ultrasonography (TCD) have been reported. The pulsality index (PI) method is implemented through calculation based on the *Q*_CBFv_, and is highly correlated with the ICP^9,10^. However, there is still debate as to whether the PI method is clinically applicable^11–13^.

In many proposed methods, a mathematical model is used to estimate ICP. Hemo-and hydro-dynamic models concerning the cerebrospinal fluid (CSF) flow dynamics were previously proposed by Ursino and Lodi, with the interaction between the cerebral blood volume, cerebral autoregulation, and ICP then being confirmed by the simple Ursino model^14–16^. Although this model describes cerebral autoregulation effectively, it is difficult to estimate the ICP because complex mathematical formulas are involved. Therefore, to estimate ICP, Kashif et al.^17^ have incorporated the *Q*_CBFv_ and *P*_a_ in a physiological model of cerebrovascular dynamics. This approach not only reflects the autoregulation mechanism, but also exhibits good performance as regards ICP estimation. However, it is inappropriate to replace the *Q*_CBF_ with *Q*_CBFv_ (the mean *Q*_CBFv_ is 80 cm/s and *Q*_CBF_ is 11.67 ml/s). Moreover, this method is unsuitable for detection of suddenly changing ICP levels, as it employs a long time window (60 s) for ICP estimation.

To overcome the limitations of the above methods, we previously proposed a simplified intracranial hemo-and hydro-dynamics model incorporating only resistance (*R*) terms and using *Q*_CBFv_ and *P*_a_, called the simple resistance (SR) model^18^. However, although the SR model has the advantage of detecting sudden changes in ICP, it has not been established for cerebral autoregulation^18^.

Therefore, in this paper, we propose a novel state-space model-based method to estimate ICP considering cerebral autoregulation in real time, called direct current (DC)-ICP. Non-invasively estimated ICP, which is calculated using the DC-ICP state-space model, and invasively measured ICP are compared using clinical data acquired from patients in a neurointensive care unit (NCU).

## Results

The mean and standard deviation of the invasive ICP and *Q*_CBFv_ values were 19.55 ± (mmHg) and 49.45 ± 21.83 (cm/s), respectively. The invasive ICP waveform was recorded using an intraventricular probe (see Methods). Each patient’s *P*_a_ waveform was also recorded simultaneously through arterial catheterization, and the *Q*_CBFv_ was recorded using TCD of the middle cerebral artery (MCA) (Fig. 1). The intracranial states, i.e., *Q*_CBF_, *P*_c_, and arterial resistance (*R*_a_), were estimated using measured variables, i.e., the *Q*_CBFv_ and *P*_a_ (Fig. 1).

**Figure 1.**
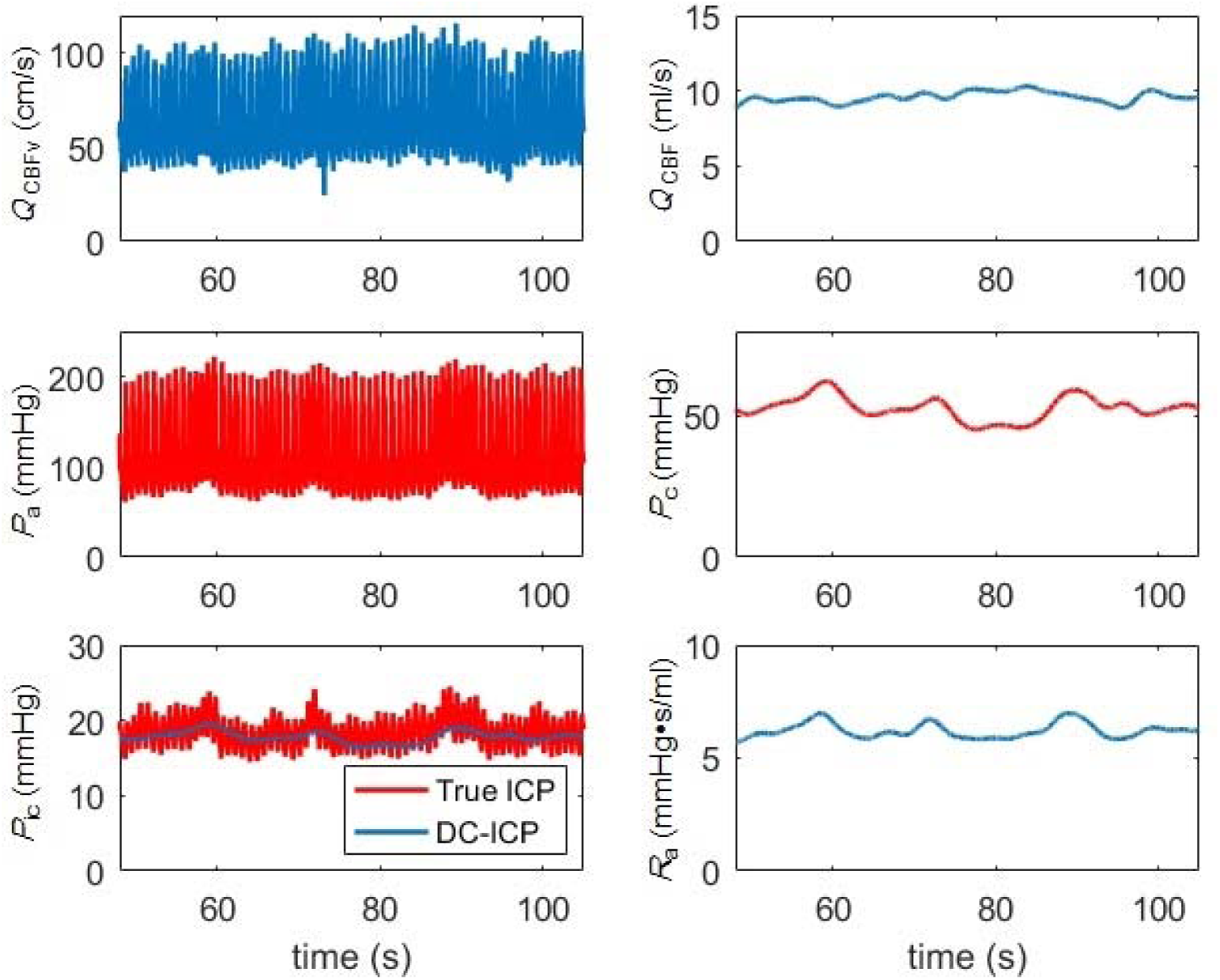
Measured and estimated intracranial state-space variables. The left column shows the measured *Q*_CBFv_ of the middle cerebral artery, the arterial pressure from the radial artery (*P*_a_), and the intracranial pressure (ICP). The right column shows the estimated cerebral blood flow (*Q*_CBF_) of the middle cerebral artery, the intracranial capillary pressure (*P*_c_), and the cerebro-vascular resistance (*R*_a_).

Figure 2 shows a comparison of the true ICP and the non-invasively estimated ICPs given by the different methods. The PI method^9,10^ cannot track changes in the actual ICP. Kashif et al.’s method^17^ reflects the true ICP trend well, but is sometimes unstable. Our DC-ICP method not only tracks the true ICP changes, but also exhibits excellent stability (Fig. 2).

**Figure 2.**
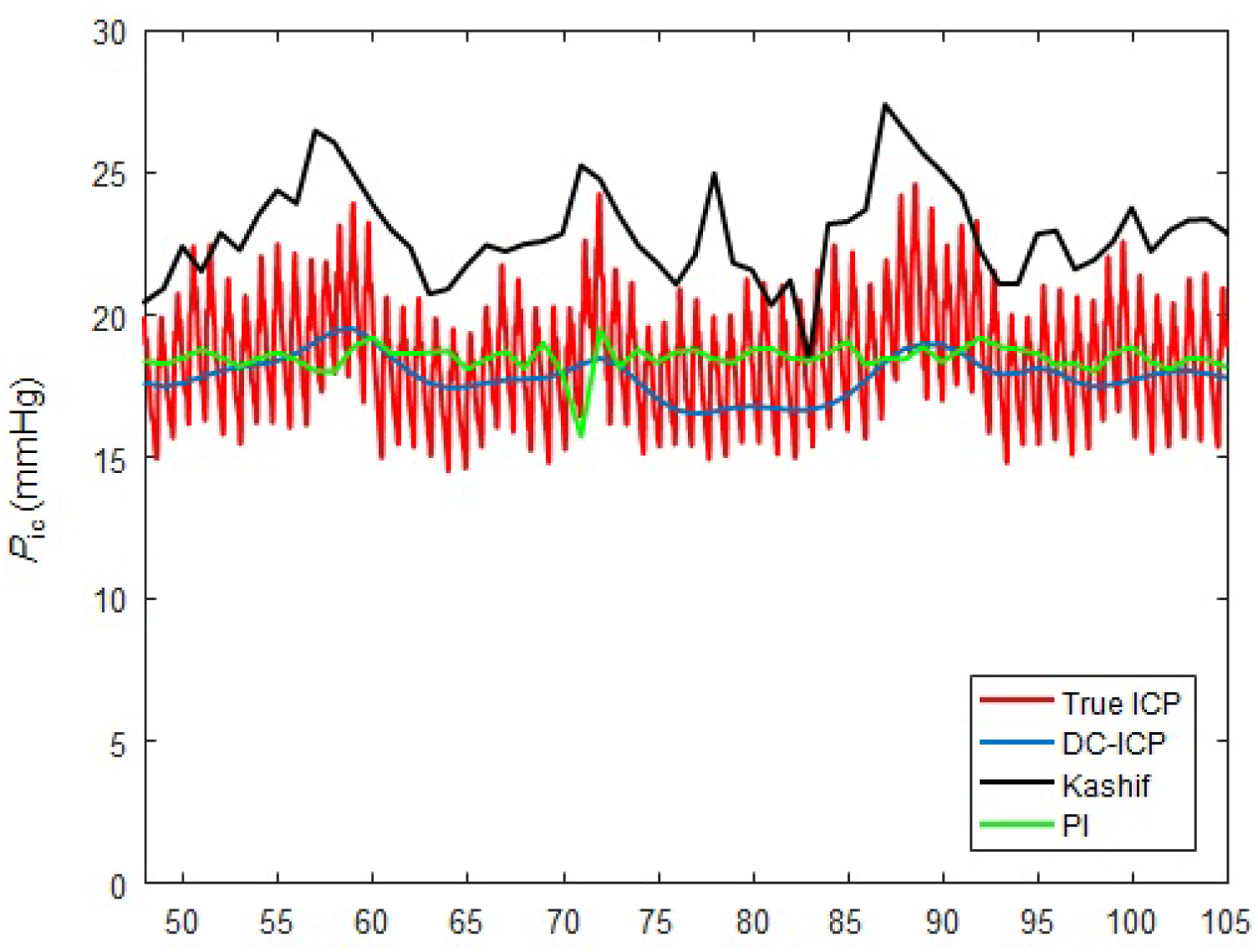
Comparison of true ICP with non-invasively estimated ICPs. The true ICP is indicated by a red line, while the results given by the pulsality index (PI)^9,10^, Kashif et al.^17^, and DC-ICP methods are indicated by green, black, and blue lines, respectively.

In Fig. 3, the bias ± standard error values of the DC-ICP, Kashif et al., and PI methods are 0.21 ± 3.52, −1.36 ± 4.57, and 0.00 ± 4.93 (mmHg), respectively. The DC-ICP exhibits not only superior precision to Kashif et al.’s method, but also the best accuracy according to the Bland-Altman plot. The inter-subject Pearson correlation coefficient between observers was 0.70.

**Figure 3.**
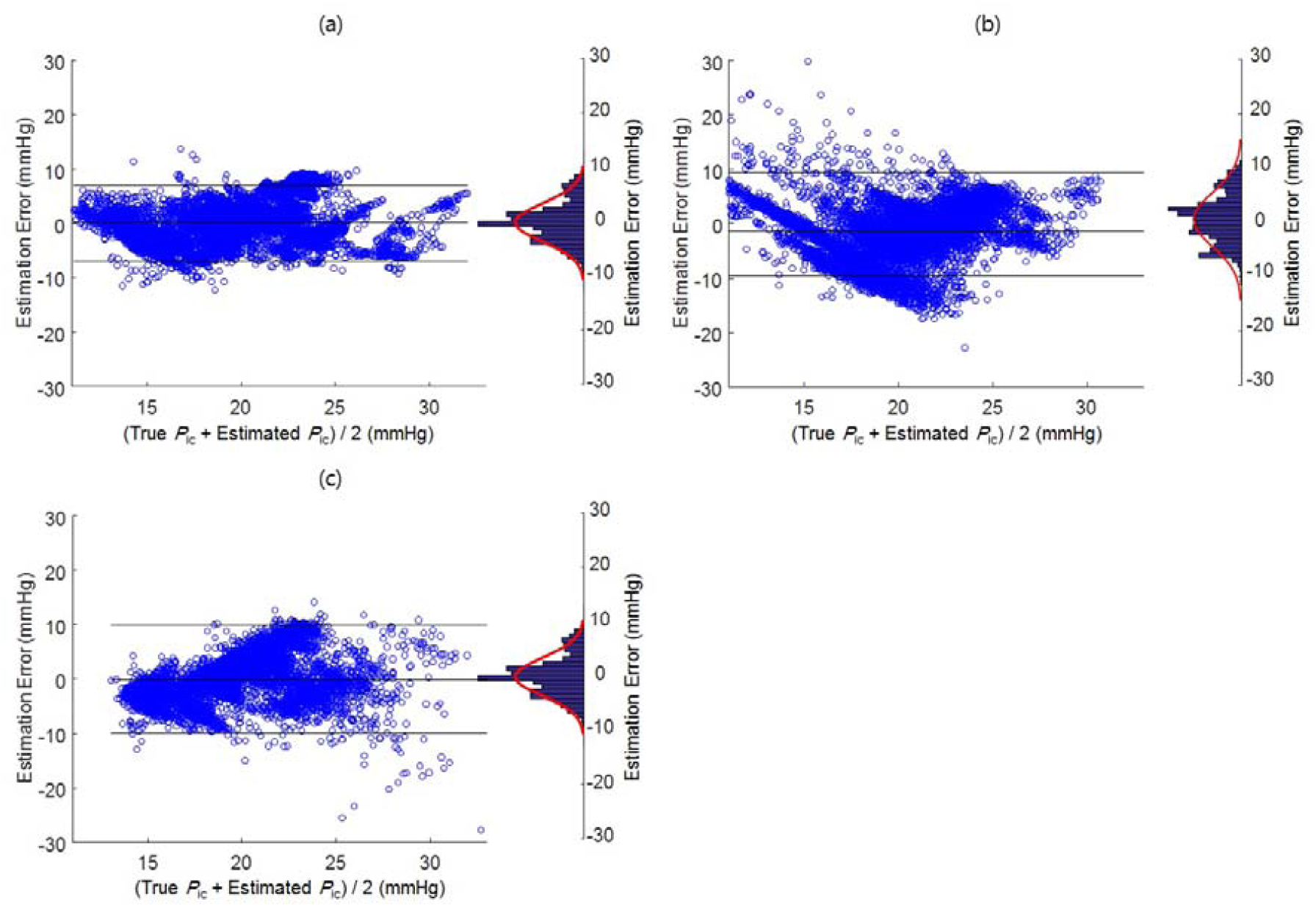
Bland-Altman plots for overall ICP estimation performance: **a**−**c** DC-ICP, Kashif et al.^17^ and PI^9,10^ methods, respectively. True and estimated ICPs over 15,256 nonoverlapping time windows of 1 s from 10 patient records are shown. The bias ± standard error values given by the DC-ICP, Kashif et al., and PI methods are 0.21 ± 3.52, −1.36 ± 4.57, and 0.00 ± 4.93 (mmHg), respectively.

Increased ICP (IICP) is clinically important, and IICP screening is the purpose of ICP monitoring. Therefore, the authors calculated the areas under curve (AUCs) of the receiver operating characteristic (ROC) curves for prediction of ICP ≥ 20 mmHg for the DC-ICP, Kashif et al., and PI methods, obtaining 0.816, 0.634, and 0.624, respectively (Fig. 4). The mean absolute error (MAE) was also calculated. The median MAEs of the DC-ICP, Kashif et al., and PI methods were 2.1612, 2.809, and 2.6693, respectively (Fig. 5).

**Figure 4.**
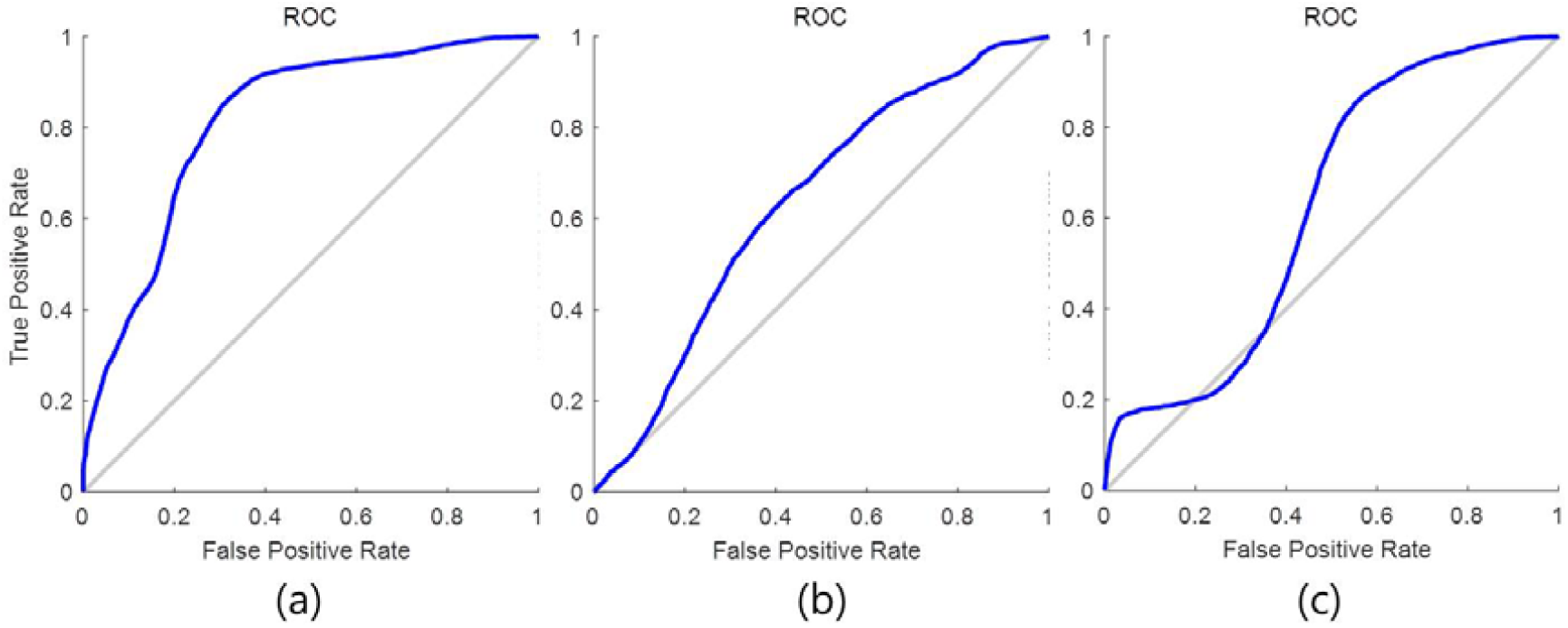
Receiver operating characteristic (ROC) curves for discrimination of elevated intracranial pressure (ICP): **a**−**c** DC-ICP, Kashif et al.^17^, and PI^9,10^ methods, respectively. The sensitivity, specificity, and area under curve (AUC) results for ICP ≥ 20 mmHg are presented. The diagonal line shows random chance (0.50). The AUCs of the ROCs for prediction of ICP ≥ 20 mmHg for the DC-ICP, Kashif et al., and PI methods are 0.816, 0.634, and 0.624, respectively.

**Figure 5.**
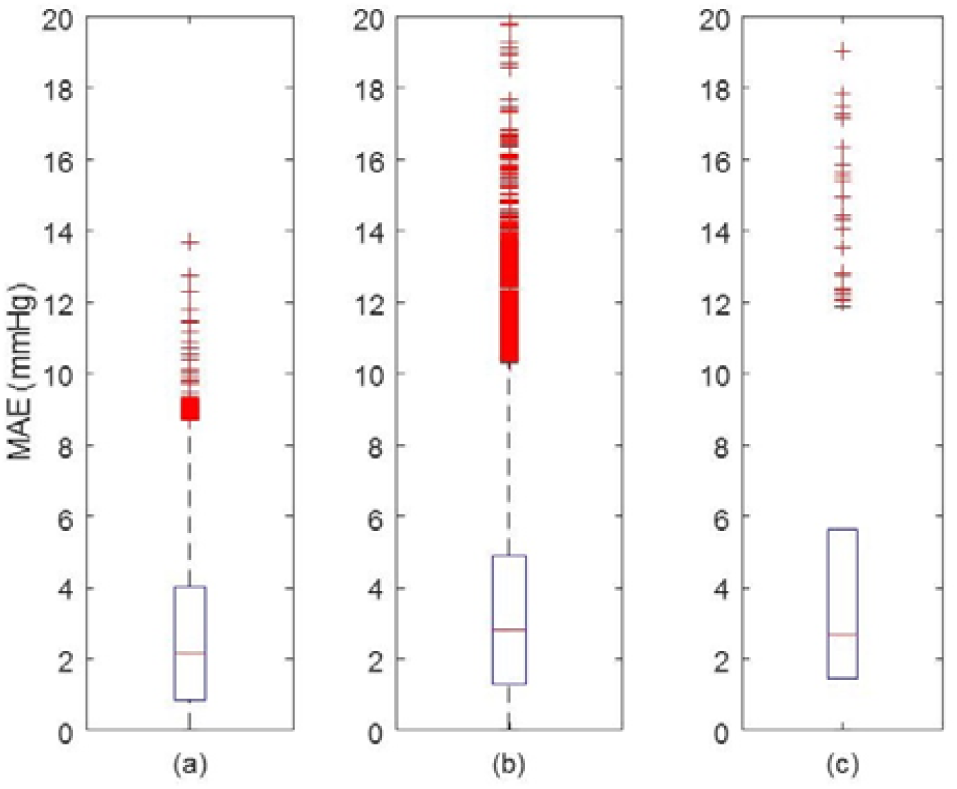
Mean absolute error (MAE) for estimated ICP: **a**−**c** DC-ICP, Kashif et al.^17^, and PI^9,10^ methods, respectively. The median MAE values for the DC-ICP, Kashif et al., and PI methods are 2.1612, 2.809, and 2.6693, respectively.

## Discussion

In this study, we suggest a novel algorithm for model-based non-invasive estimation of intracranial states such as *P*_c_, *Q*_CBF_, and the ICP. Considering figure 2, The PI method cannot trace the changes in the true ICP. Kashif et al.’s method reflects the true ICP trend well, but with low precision. Our DC-ICP method not only traces the true ICP changes, but also exhibits good precision. Our results show that the non-invasively measured ICP estimated using *Q*_CBFv_ and *P*_a_ has a statistically significant correlation with the invasively measured ICP. Moreover, the observed detection of IICP (ICP ≥ 20 mmHg) is promising for clinical application (correlation coefficient: 0.70; AUC for prediction of ICP ≥ 20 mmHg: 0.81).

Because of its non-invasive nature, TCD has been widely used to detect changes in cerebral blood flow of patients in NCU, to assess cerebral vasospasms, circulatory disorders, and other brain injuries^9,19–22^. A *Q*_CBFv_ can be measured using TCD, from which systolic and end-diastolic flow rates can be obtained. Routinely, the mean flow velocity, resistance index, and PI have been derived from the measured waveform result.

Various studies have been performed to noninvasively estimate ICP using these TCD measurements^9,11,12,23–25^. Continuous decreases in the mean flow velocity and end diastolic velocity with rising ICP, as well as a rise in the resistance index, have been reported^23,24^. Some researchers have reported that the PI is also associated with the ICP^9,25^. However, in patients undergoing lumbar shunt insertion, lumbar infusion tests performed to determine changes in ICP revealed no relationship between the PI and ICP^11^. In addition, the PI tends to be less correlated with the ICP when the ICP rises^12^. As suddenly increasing ICP does not affect the *Q*_CBFv_ waveform, the PI-based approach cannot detect sudden development of intracranial hypertension and does not reflect cerebral autoregulation. In our study, we observed a trend consistent with these previous findings. For clinical data, ICP estimation using PI was compared with the actual ICP. However, the ability to predict increased ICP was poor (correlation coefficient: 0.43; AUC for prediction of ICP ≥ 20 mmHg: 0.55). Therefore, use of PI to estimate ICP is inaccurate.

Recently, a method of estimating ICP by measuring the arterial pulsation of the intra-and extra-cranial segments of the ophthalmic artery using two-depth Doppler was introduced^26,27^. The correlation coefficient between invasive and non-invasive ICP measurements was 0.74. The sensitivity, specificity, and AUC for ICP > 20 mmHg were 0.72, 0.77, and 0.71, respectively. The disadvantage of this method is that it is difficult to monitor the ICP continuously for a long time. Further, the measurement is unreliable in cases involving eye problems such as glaucoma, or compartment status with various causes. In addition, for this type of ICP measurement, the clinician must purchase specific equipment and cannot use conventional TCD.

To resolve the continuous monitoring problem, model-based ICP estimation methods have been developed and examined^14,17,28–35^. Hu et al. first suggested the model-based ICP estimation approach based on the Ursino model, and estimated various intracranial states including ICP from noninvasive measurements (*P*_a_ and *Q*_CBFv_)^32^. To suppress the instability caused by the model complexity, they employed a Kalman filter combined with quadratic programming. However, it is difficult to apply this approach to ICP estimation in an actual clinical scenario, because the Ursino model contains too many parameters (such as the arterial compliance, *R*, and elastance coefficient), and the values cannot be determined adaptively. Further, the estimates are determined by these model parameters. To solve the problem of model parameter selection, Kashif et al.^17^ developed an innovative algorithm based on a simplified model. The correlation between the non-invasive ICP and invasive ICP measurements was an correlation coefficient of 0.90, sensitivity of 83%, specificity of 70, and an AUC of 0.83 for prediction of ICP ≥ 20 mmHg. Kashif et al. estimated the ICP using the radial artery pressure and *Q*_CBFv_ only, through model simplification. This method not only estimates the ICP through very simple equations, but also constitutes a very innovative advance in terms of adaptive learning of model parameters such as resistance and compliance of artery. However, the adaptation algorithm depends on the waveform morphology instead of the physiological mechanism. In that context, *Q*_CBF_ is substituted for *Q*_CBFv_, which is inappropriate (the mean *Q*_CBFv_ is 80 cm/s and *Q*_CBF_ is 11.67 ml/s). The most prominent disadvantage is that it is difficult to track clinically rapid ICP changes using this approach, because a long time window is essential (60 s).

In a previous study, we proposed a new algorithm, the SR model, to overcome this problem^18^. As the ICP to be estimated is the DC level of the pressure and not the AC fluctuation, we removed the AC component of the measured signals (*P*_a_, *Q*_CBFv_) in the signal preprocessing step through the low-pass filter. Consequently, the *C* term of the Ursino model was removed. Then, the ICP could be estimated using a simple equation with good performance. However, a limit existed in that simulation-based validation is performed without validation through actual clinical data available at that time. In addition, there is no adaptation algorithm for *R*_a_ selection, unlike the Kashif et al. method, which is controlled through autoregulation^36^. Therefore, in this study, we devised an algorithm that reflects the autoregulation model to automatically estimate *R*_a_. This algorithm was then validated using clinical data.

The proposed DC-ICP estimation method has several advantages. First, several intracranial states (*Q*_CBF_, *R*_a_, and ICP) as well as the ICP can be estimated using the newly proposed model, employing an unscented Kalman filter (see Methods). The estimated *Q*_CBF_, *R*_a_, and ICP can help clinically monitor patient status. Second, most non-invasive methods require calibration for ICP estimation; however, the method proposed in this study allows the clinician to estimate the ICP without additional calibration. Third, Kashif et al.’s method does not reflect abrupt changes in ICP, because data measured over 60-s periods are used. However, using our proposed algorithm, continuous and real-time (latency: 1 s) ICP monitoring is possible. Finally, *Q*_CBF_ can be estimated in a realistic manner. Previous model-based ICP estimation approaches substituted *Q*_CBFv_ for *Q*_CBF_ with or without proportional regulation (*Q*_CBF_ = *k* × *Q*_CBFv_, where *k* is a constant). Hence, the proposed estimation algorithm is more accurate than previous approaches.

Our work has certain limitations. First, our proposed algorithm was not validated for general neurological systems, because of the need for invasive ICP monitoring in the patients included in our study. However, patients who require ICP monitoring have similar indication to those considered in this study; thus, there is a clinically relevant aspect. Second, only 10 patient cases were considered for validation of the proposed method; this number is insufficient compared with previous studies. To overcome the low sample-size problem, our algorithm was constructed with a 1-s epoch and analyzed over a total of 15329 time points. Third, *Q*_CBFv_ and *P*_a_ should be measured simultaneously for ICP estimation. However, most patients who require our algorithm are A-line monitored, and *Q*_CBFv_ is a commonly measured value in NCU employing TCD. Fourth, as the venous sinus pressure is set to constant, errors may occur depending on the volumetric status of the patient. In this work, we considered the intake/output balance to maintain the patients’ euvolemic status as much as possible.

## Outlook

Invasive ICP monitoring is fundamentally important and must be performed in patients who are eligible for external ventricular drain insertion, because ICP regulation via CSF drainage as well as monitoring can be applied. However, there are certain clinical situations in which invasive ICP monitoring is not possible. Therefore, our ICP estimation algorithm based on the intracranial state-space model is expected to provide valuable information under clinical conditions in specific situations.

## METHODS

### Data Acquisition

This is a two-center (Gangnam Severance Hospital and Cheju Halla Hospital), prospective observational study conducted from 6 July 2017 to 1 March 2018. 10 patients with a mean age of 57 years were admitted to the respective NCUs with TBIs and required invasive neurosurgical monitoring (Table 1). In total, 15329 s of data were collected, with 1 s being defined as an epoch for estimation. Based on the correlation between our model and that of Kashif et al.^17^, we constructed 255 epochs based on 60 s. All patients were aged 18−70 years and had suffered a TBI with intracranial hemorrhage (ICH). They required clinically invasive ICP monitoring and TCD. For each patient, the following characteristics were collected: GCS at admission, age, sex, height, weight, brain injury mechanism, and Glasgow Outcome Scale (GOS) score at discharge. The patients’ representatives gave informed consent. The institutional review boards of the Yonsei University Gangnam Severance Hospital, Cheju Halla General Hospital, and Gwangju Institute of Science and Technology approved our study, respectively.

**Table 1.** Patient Demographics

| Case No. | Sex | Age (years) | Height (cm) | Weight (kg) | Initial GCS (score) | Pathology | GOS at discharge (score) | Comorbidities |
| --- | --- | --- | --- | --- | --- | --- | --- | --- |
| 1 | F | 47 | 160 | 57 | 5 (E1V1M3) | ICH | 2 | HTN, DM |
| 2 | M | 86 | 165 | 70 | 4 (E1V1M2) | SAH | 2 | none |
| 3 | F | 50 | 162 | 55 | 3 (E1V1M1) | SAH | 4 | Hypothyroidism |
| 4 | F | 49 | 153 | 60 | 9 (E2V3M4) | SAH | 4 | HTN |
| 5 | F | 54 | 160 | 50 | 13 (E3V4M6) | SAH | 5 | none |
| 6 | M | 50 | 167 | 73 | 5 (E1V1M3) | ICH | 1 | none |
| 7 | M | 52 | 170 | 90 | 8 (E2V2M4) | SAH | 5 | HTN |
| 8 | F | 54 | 160 | 63 | 3 (E1V1M1) | ICH | 3 | HTN |
| 9 | M | 74 | 170 | 60 | 9 (E3V1M5) | ICH | 3 | HTN |
| 10 | M | 54 | 158 | 60 | 10 (E3V3M4) | SAH | 2 | HTN |

Initial GCS(E: eye movement, V: verbal communication, M: motor response), GOS: glasgow outcome scale, ICH: intracranial hemorrhage, SAH: subarachnoid hemorrhage, HTN: hypertension, DM: diabetes mellitus

### Intracranial pressure and cerebral blood flow velocity measurement protocol

The ICP was measured invasively via a catheter inserted into the brain ventricles and connected to an external pressure transducer and drainage system (Codman, Johnson & Johnson, Raynham, Massachusetts, US). The ICP was recorded by an assisting nurse. *Q*_CBFv_ measurement was performed using TCD. The examiner performing this measurement was blind to the ICP monitoring results. TCD was performed on the MCA through a temporal window using a traditional 2-MHz transducer (Ez Dop; DWL, Singen, Germany) with the head elevated to 30° (Fig. 6).

**Figure 6.**
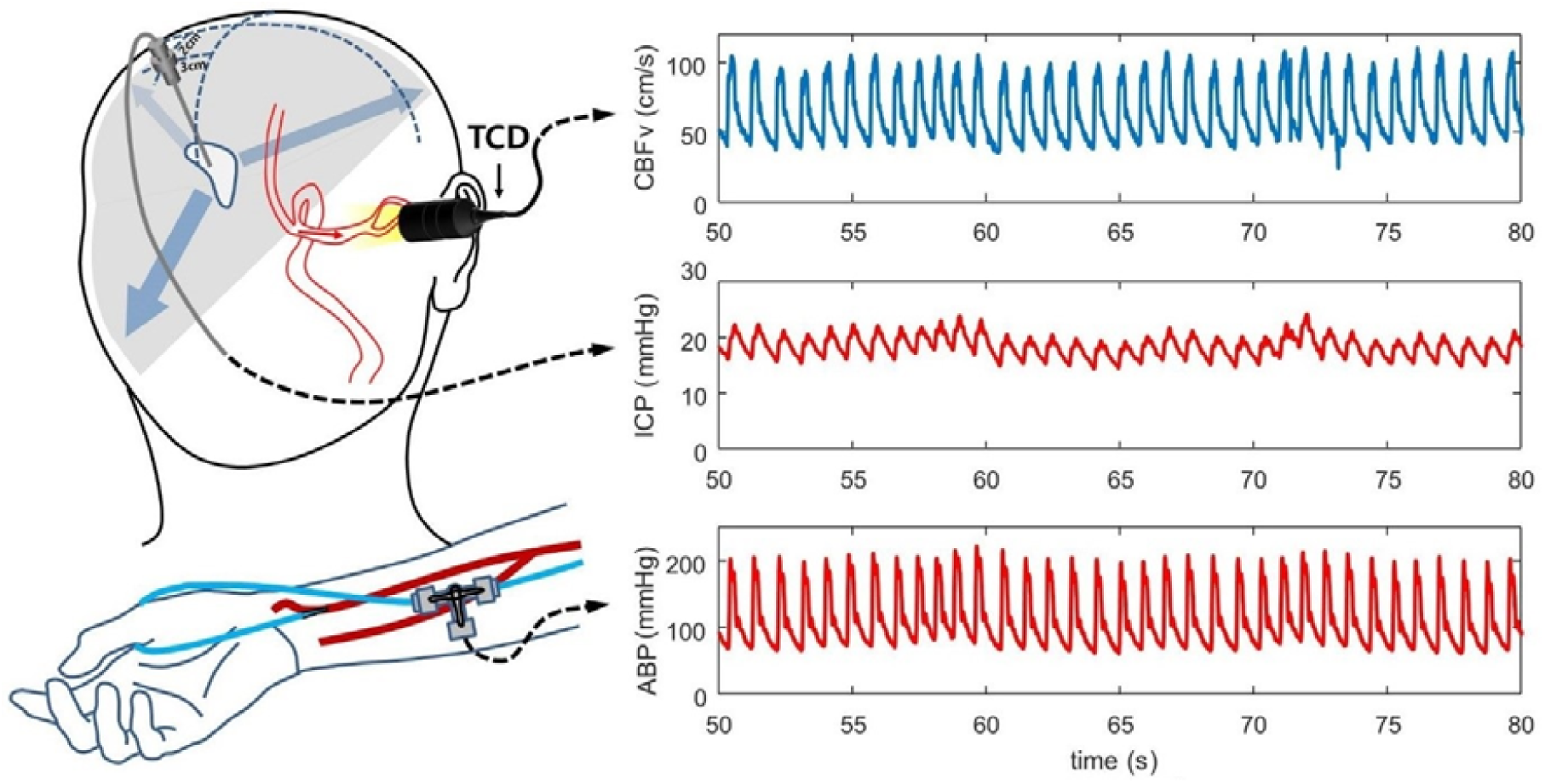
Measured arterial blood pressure (*P*_a_), cerebral blood flow velocity (*Q*_CBFv_), and intracranial pressure (ICP). The *Q*_CBFv_ was measured using transcranial Doppler sonography (TCD) at the middle cerebral artery. The ICP was acquired using an intraventricular monitoring device. The *P*_a_ was obtained from the radial-artery A-line.

### Existing models modified from original Ursino model

### Simplification of original Ursino model: SR model

The original Ursino model^14–16^ is an intracranial hemo-and hydro-dynamic model expressed using both resistance (*R*) and compliance (*C*) terms. As the *C* term has infinite impedance with DC input, all *C* terms of the original Ursino model can be cancelled when the input has a DC trend only. This simplified result is the SR model^18^ (Fig. 7). The ICP can be estimated using simple equations based on the SR model, as follows:

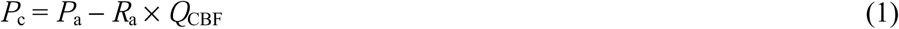

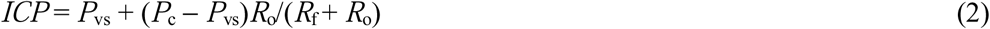

where *P*_a_, *P*_c_, ICP, and *P*_vs_ are the arterial, capillary, intracranial, and sinus venous pressures, respectively; *Q*_CBF_ represents the CBF; and *R*_a_, *R*_o_, and *R*_f_ are the mean arterial, CSF formation, and CSF outflow resistances, respectively. All pressures and flow have DC trends, and a DC trend input (*P*_a_ and *Q*_CBF_) can be simply implemented through low pass filtering (∼0.5 Hz).

**Figure 7.**
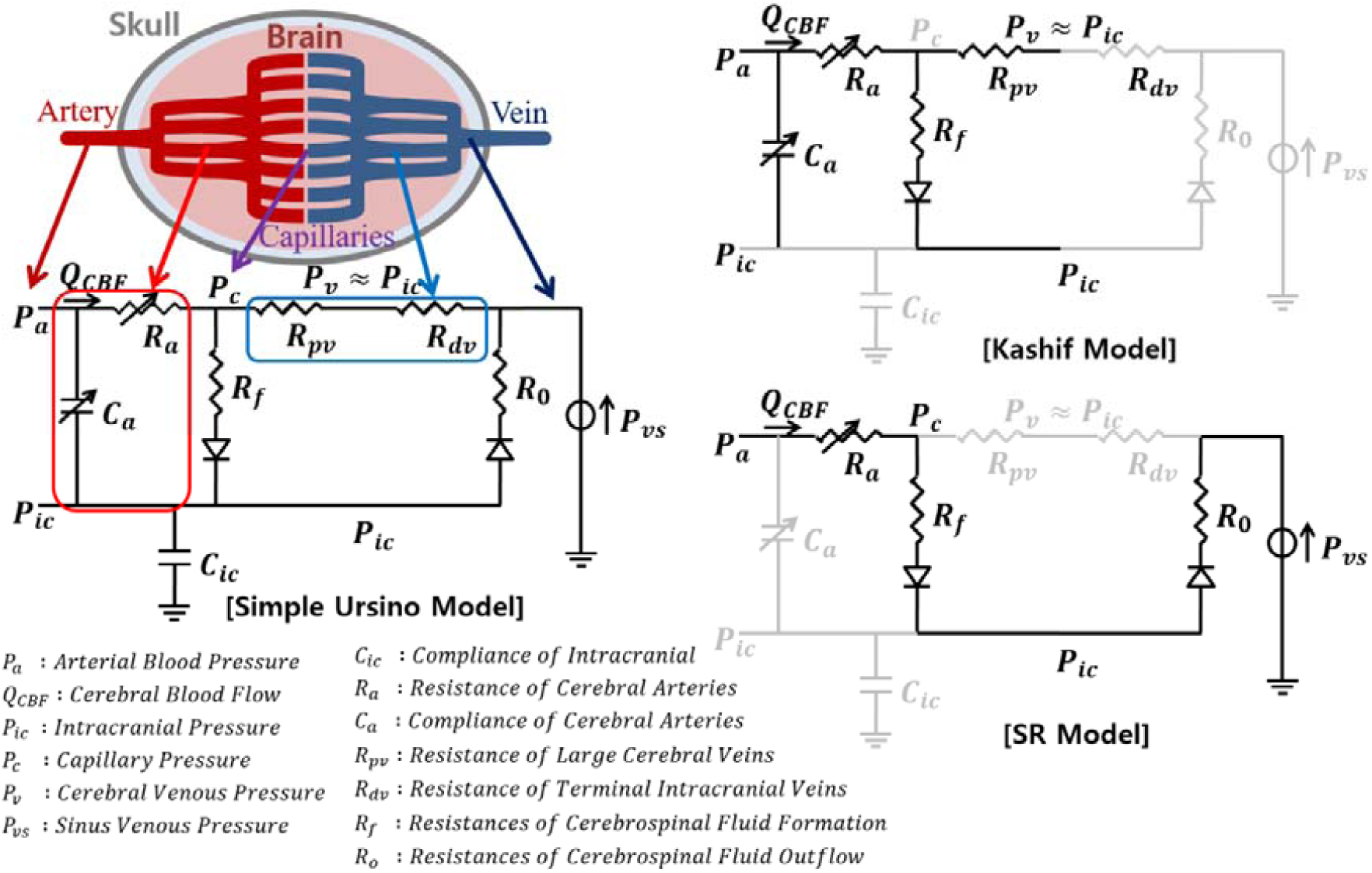
Existing models, which are modifications of the original Ursino model. The original Ursino model is an intracranial hemo-and hydro-dynamic model incorporating both resistance (*R*) and compliance (*C*) terms. The Kashif et al. and SR models can be regarded as simplified models of the original Ursino model.

### Kashif et al. model

The Kashif et al. model^17^ has two components (Fig. 7), *C* and *R*, and their estimates (Ĉ and 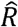) are calculated as follows:

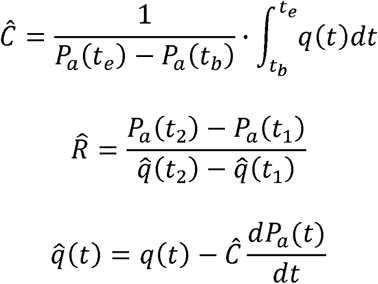

where *t*_b_ and *t*_e_ are the time instants of the beginning and end of the systolic upstroke in *P*_a_(*t*); and *t*_1_ and *t*_2_ are time instants near the local minimum and maximum of *P*_a_(*t*). The noninvasive ICP is given by

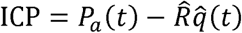

### State-Space Model-based ICP Estimation

### Cerebral auto-regulation model: arterial resistance and its cross-sectional area

Although *R*_a_ is regulated, it is fixed at 6 (mmHg s/ml) in the SR model^18^. Thus, an autoregulation model for *R*_a_ must be applied. An autoregulation device (ARD)^37^ is an arterial resistance model for autoregulation, which is expressed as follows:

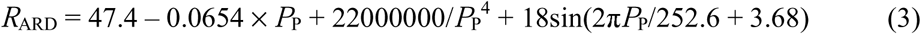

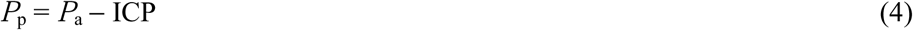

where *P*_P_ is the cerebral perfusion pressure. Here, *R*_a_ corresponds to part of *R*_ARD_ and their units are different (R_ARD_: mmHg·0.1 kg·60 s/mL, R_a_: mmHg·s/mL). Therefore, to match the unit scales of *R*_ARD_ and *R*_a_, we approximate *R*_a_ by dividing *R*_ARD_ by 6.

Knowledge of the arterial cross-sectional area *A*_a_ is important, because it is related to the unobservable *Q*_CBF_, where

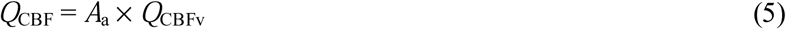

Note that *A*_a_ can be derived from the given *R*_a_ using Poiseuille’s law:

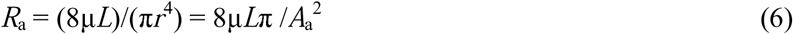

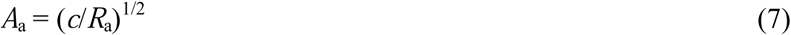

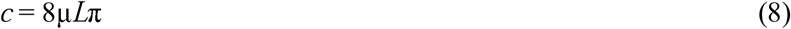

where μ, *L*, and *r* represent the viscosity, length, and radius of the vessel, respectively. In this study, we assumed constant μ and *L* for simplicity. Note that the *A*_a_ of equation (5) is not precisely identical to that of equation (7), because the *Q*_CBFv_ of equation (5) is measured from just one location on the arteries (e.g., the MCA), whereas the *A*_a_ of equation (7) represents the overall arterial cross-sectional area. However, as these two values are correlated, we assumed that the MCA cross-sectional area was equivalent to that for the overall arteries in this work.

### State estimation using unscented Kalman filter (UKF): DC-ICP algorithm

From equations (1)−(8), we constructed a state-space model to estimate the intracranial states, which is expressed as follows:

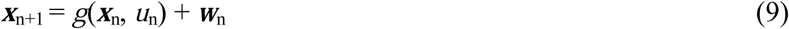

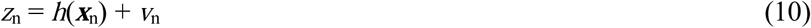

where ***x***_n_= [*Q*_CBF_ *P*_c_ ICP *R*_a_]^T^, *u*_n_ = *P*_a_, and *z*_n_ = *Q*_CBFv_. Here, ***x***_n_, *u*_n_, *z*_n_, ***w***_n_, and *v*_n_ indicate the state, input, observation, system noise, and measurement noise, respectively. Note that *<u>g</u>*(**x**_n_, u_n_) is composed of equations (1)−(4), and *h*(***x***_n_) comprises equations (5) and (7).

To estimate the intracranial state ***x***_n_ from the given state-space model, we adopted an optimal state estimation technique, which estimates a hidden state based on a state-space model^38^. In this study, we utilized an unscented Kalman filter (UKF), as this technique provides one of best solutions for a nonlinear state estimation. Note that we refer to the proposed method as the DC-ICP because it is focused on the ICP direct current (DC).

### Algorithm Validation in TBI Patients

To validate the proposed DC-ICP algorithm, we compared it with previously reported methods; namely, the Kashif et al.^17^ and PI^9,10^ methods. The PI method estimates the ICP from linear regression of the PI as follows:

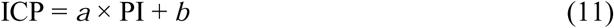

where *a* and *b* are regression coefficients. As estimates given by the PI method have several estimation biases, we performed additional calibration with offset value.

### Statistical analysis

ICP values measured using the invasive manner were compared with non-invasive ICP values estimated by our proposed, the Kashif et al.^17^ and PI^9,10^ methods. Bland-Altman analysis was performed to determine the inter-method agreement. The correlation between methods was assessed by considering the Pearson correlation coefficient. Receiver operating characteristic (ROC) curves were constructed to determine the sensitivity, specificity, and area under the curve (AUC) of the estimated values compared with the measured values for elevated ICP (≥20 mmHg). Statistical significance was defined as *P* < 0.05. Statistical analysis was performed using SPSS version 22 (IBM, Armonk, New York, USA) and MATLAB version R2016a (MathWorks, Natick, Massachusetts, USA).

## Acknowledgments

This work was supported by the GIST Research Institute (GRI) in 2018, and by the National Research Foundation of Korea (NRF), grant funded by the Korean government (MSIP) (2017R1A2B4009068).

## Author contributions

Boreom Lee conceived the idea of the project. C. Park, S. Ryu and S. Lee organized the project. C. Park, S. Ryu and B. Jung collected human data. S. Lee, C. Hong and Y. Kim performed surgeries, recorded patient neurologic status. C. Park and S. Ryu performed the analysis of the acquired data. C. Park and S. Ryu wrote the manuscript. B. Lee led the project.

